# Metaxa: A Transformer-Based Deep Learning Model for Taxonomic Classification of Long Nanopore Reads

**DOI:** 10.64898/2026.04.20.719780

**Authors:** Krešimir Friganović, Dominik Stanojević, Poshen B. Chen, Mile Šikić

**Author notes:** Corresponding author: Mile Šikić. Contributed equally (Krešimir Friganović, Dominik Stanojević).

## Abstract

A significant fraction of the microbial diversity remains unclassified, hindering our understanding of microbial roles in health and ecosystems. State-of-the-art methods like Kraken 2 perform well for taxa that are present in the database. However, their accuracy drops significantly when classifying taxa that are not included. While deep learning has advanced many fields, its applications in metagenomics remain limited, and its full potential has yet to be realized. Here, we present Metaxa, a transformer-based deep learning model designed for the taxonomic classification of long-read Nanopore sequences. Metaxa leverages the sequential context of Nanopore reads, enabling robust classification beyond fixed k-mer profiles. Our results show that Metaxa matches Kraken 2 on in-sample data at both the species and genus levels, and significantly outperforms both Kraken 2 and MetageNN at the genus level on out-of-sample datasets where the species genome is absent from the reference database but a different species from the same genus is present. Furthermore, Metaxa demonstrates strong generalization across different Nanopore chemistries (R9.4.1 and R10.4.1). This work highlights the potential of deep learning models to improve metagenomic classification accuracy, especially in complex or underexplored environments where traditional tools fall short.

## 1 Introduction

Despite intense study of the microbial communities, a significant fraction of microbes remain unknown or unclassified. These “microbial dark matter” species are not represented in current reference databases or cultures [1][2], yet they may play crucial roles in human health and ecosystems.

Even in the well-studied human microbiome, a portion of genetic material cannot be assigned to any known organism. Studies estimate that on average about 2% of sequences in human microbiome samples are taxonomically unknown, and in certain environments (such as skin or oral microbiomes) this can rise to 25% [3]

Unidentified microbes could be the hidden factors in medical conditions or the sources of new therapeutic opportunities. For example, the discovery of new gut species enriched in patients with inflammatory bowel disease or other gastrointestinal disorders suggests that previously unknown organisms might be contributing to disease progression or symptoms [4]. If we do not know these species exist, we cannot link them to disease or target them with treatments. On the flip side, unknown members of the microbiome might be beneficial – producing vitamins, anti-inflammatory compounds, or other metabolites crucial to our well-being. By identifying them, we could harness their activities.

The improvement in database curation and the building of new tools increase our ability to detect and quantify taxa in the samples. Metagenomics has revolutionized microbial research by enabling direct sequencing of genetic material from diverse environments, providing insights into microbial diversity, ecosystem dynamics, and human health. Nanopore sequencing allows the generation of long reads, offering unique opportunities for metagenomic taxonomic classification.

The metagenomic taxonomic classification aims to determine the composition of a microbial community by assigning sequencing reads to taxonomic groups. Traditional approaches can be broadly categorized into k-mer-based methods, mapping-based strategies, and protein-level similarity searches [5]. Tools such as Kraken [6], Kraken 2 [7], and Centrifuge [8] employ k-mer-based strategies, while others, such as MetaPhlAn [9], rely on marker genes. These tools are primarily designed for short-read sequencing, and no definitive “best” classifier has emerged [10]. Moreover, k-mer-based methods like Kraken 2 struggle with unsequenced species or incomplete reference genomes, often leading to incorrect taxonomic assignments or labels of “unknown” taxa [11]. Mapping-based tools like MetaMaps [12] align reads to reference genomes and estimate taxonomic abundances based on mapping quality and genomic coverage, making them suitable for long-read sequencing technologies. However, they can be computationally intensive and slower than k-mer-based methods, especially when applied to large reference databases, and may still produce incorrect classifications in the presence of highly similar genomes. Therefore, there is a need for new approaches that might close this gap.

Recent advances in artificial intelligence (AI), particularly in natural language processing (NLP) [13][14][15], have introduced new possibilities for biological sequence analysis. Deep learning models, such as transformers, have demonstrated strong performance in sequence-based tasks, including gene expression prediction [16], splice site recognition [17], and epigenomics [18][19]. Recognizing their potential for metagenomics, models such as DeepMicrobes [20], BERTax [21] and MetaTransformer [22] have been developed for short-read taxonomic classification. While promising and potentially useful for classifying unknown taxa at higher taxonomic ranks, these models generally perform worse than traditional methods, especially at the species level.

The recently developed MetageNN [23] is specifically designed for long Nanopore reads and performs well at the genus level when classifying out-of-sample species compared to algorithmic approaches. However, MetageNN relies on short k-mer profiles for taxonomic classification. Although this approach helps mitigate the impact of ONT read errors, both during training (where reference genomes are assumed to be error-free) and inference (since shorter k-mers are less likely to contain sequencing errors), it neglects the valuable sequential information provided by long-read sequencing.

We developed Metaxa, a transformer-based deep learning model tailored for the taxonomic classification of long nanopore reads. Metaxa builds on the strengths of transformer architectures to encode dependencies in erroneous DNA sequences. Our evaluation demonstrates that Metaxa performs comparable to Kraken 2 on in-sample datasets and significantly outperforms it on out-of-sample datasets. Moreover, we highlight its robustness across different nanopore sequencing chemistries (R9.4.1 and R10.4.1).

## 2 Materials and methods

### 2.1 Method overview

Metaxa is a deep-learning model designed to assign nucleobase sequences (i.e., reads) to species and genus taxonomy levels. The overall processing pipeline, from sequence chunk extraction to classification, is illustrated in Fig. 1. The model first encodes canonical nucleobases (‘A’, ‘C’, ‘G’, ‘T’) using one-hot encoding, generating an encoded sequence representation *X*^*L*×4^. This representation is processed through a 1D convolutional layer Conv1d(*X, d, k, s*), where *d* denotes the number of output channels, *k* denotes the kernel size and *s* denotes the stride. We set *d* = 768, *k* = 31 and *s* = 5. We selected *k* = 31 to be consistent with the minimizer length used in Kraken 2 [7]. The convolution layer effectively extracts every fifth 31-mer from the sequence, resulting in a tensor with a reduced length of 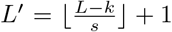. This reduction substantially decreases computational costs in the subsequent transformer encoder by mitigating the quadratic time complexity associated with self-attention, while preserving relevant information.

**Figure 1:**
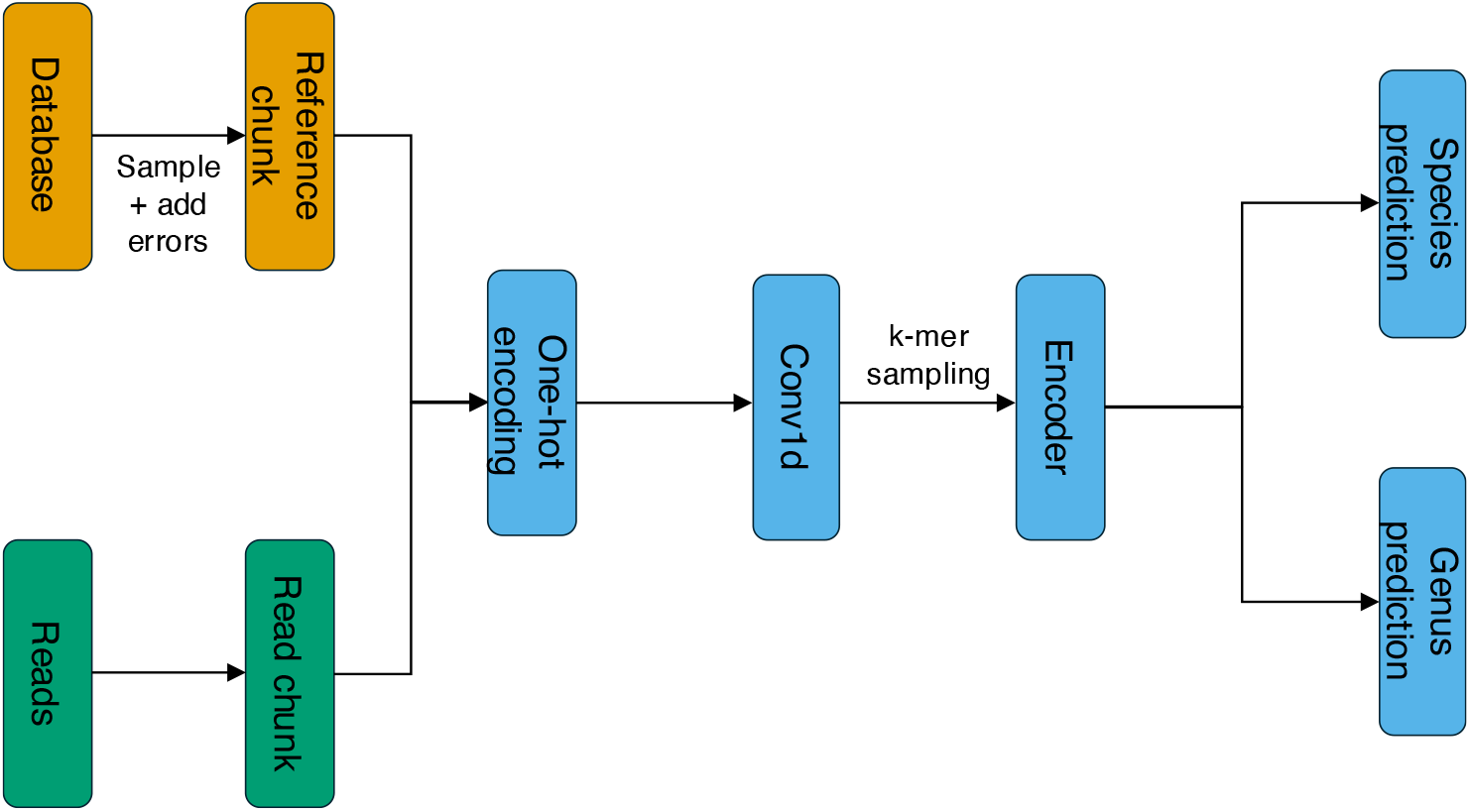
Block diagram of the Metaxa tool. Training begins with sampling a reference chunk from the database and introducing errors (orange). The sequence is then one-hot encoded, and a 1D convolution creates and samples k-mers. A Transformer encoder learns contextual representations for predicting species and genus categories. During inference, chunks are extracted from reads (green) and processed identically to training (shared steps in blue).

A classification token of size *d* is prepended to the convolution output, serving as an aggregator of information across the sequence for downstream classification tasks. The resulting tensor is passed through a transformer encoder, which generates contextualized representations for each encoded 31-mer. We use a Pre-Layer Normalization (Pre-LN) [24] transformer encoder, which stabilizes training and improves convergence by normalizing inputs before the attention and mixer layers. Our transformer encoder consists of 12 encoder layers, each with 12 attention heads, a feedforward dimension of 4*d* = 3072, and a dropout rate of 0.1. Moreover, to encode positional information, we use Rotary embeddings [25] and replace LayerNorm [26] with the more computationally efficient RMSNorm [27]. To further optimize performance, we use FlashAttention-2 [28], an algorithm designed to accelerate the attention layer in transformers. It improves speed and memory efficiency, enabling the processing of longer sequences without excessive computational resources. From the encoder’s output, the classification token, containing aggregated sequence information, is extracted and passed into separate species and genus classification heads. Each head consists of a single linear layer without a bias term, producing outputs that correspond to unnormalized probabilities. The output dimensions match the number of species and genus categories in the training database.

### 2.2 Training

During training, cross-entropy loss was used for species loss ℒ_*species*_ and genus loss ℒ_*genus*_. The total loss is calculated as the sum of these losses: ℒ_*total*_ = ℒ_*species*_ + ℒ_*genus*_. We used the AdamW optimizer with a weight decay of 1 ×10^−4^. The initial learning rate was set to 3 ×10^−4^ and decayed to 1 ×10^−6^ during training using a cosine decay schedule. Gradients were clipped if their total *ℓ*^2^-norm (computed across the individual gradient norms) exceeded 10.

The model was trained for 1 million iterations with an effective batch size of 8192, achieved by training on 8 NVIDIA A100 GPUs with a batch size of 1024 per GPU. To further reduce GPU memory usage during training, activation checkpointing was used. Validation was performed every 10,000 steps with 100 validation steps per run. Each step included 16,384 randomly sampled examples. The model with the lowest validation loss, calculated using the same criteria as the training loss, was selected as the final model.

### 2.3 Inference

At inference time, our model processes an input FASTA/Q file and generates predictions for species and genus classification for each read. Reads are divided into non-overlapping chunks of 1000 bp, and any chunk shorter than 1000 bp is discarded. For each generated chunk, the model outputs unnormalized probabilities (logits) for both species and genus classifications. The final predictions for each read are obtained using soft voting, where the probabilities for each species and genus are averaged across all chunks extracted from the read. The species and genus with the highest average probabilities are selected as predicted classifications. This approach ensures robust predictions by integrating information across all chunks of a read.

### 2.4 Training database and example generation

We based our training database on the main database described in the Meta-geNN paper [23], with several modifications to improve its quality. First, we extracted the species taxonomic IDs (taxids) for the genomes in the original database, yielding 506 unique species taxids. We then downloaded only the genomes corresponding to these taxids that were labeled as reference genomes, resulting in 504 genomes representing 502 species. Two species, *Escherichia coli* and *Sodalis praecaptivus*, had two distinct assemblies labeled as reference genomes, all of which were included. Four species (*Deinococcus soli, Rickettsia sp. MEAM1* (*Bemisia tabaci*), *Thermovirga lienii* and *Nitrosomonas stercoris*) lacked reference genome assemblies and were excluded. To enable compatibility with the evaluation datasets from ZymoBIOMICS High Molecular Weight (HMW) DNA Mock Microbial (Zymo D6322), we added reference genomes for four additional species: *Pseudomonas aeruginosa, Salmonella enterica, Enterococcus faecalis*, and *Listeria monocytogenes*. The final database consists of 508 reference genomes from 506 species across 313 genera.

Both training and validation examples are sampled from the training database and are generated dynamically during training as follows. First, a species is selected randomly, with each species having an equal probability of being chosen. If the selected species has multiple assemblies in the database, one assembly is chosen at random, again with equal probability for each assembly. Next, a contig is selected from the chosen assembly, with the selection probability proportional to the contig length, ensuring that longer contigs are more likely to be chosen. This approach ensures uniform sampling from the chosen assembly. Finally, a starting position within the chosen contig is determined uniformly at random. From this position, a 1000 bp sequence is extracted. To simulate the circular nature of the genome, bases from the beginning of the contig are appended to the sequence fragment if it is shorter than the desired length (*L* = 1000 bp), ensuring that the required length is achieved.

Once the sampled sequence is obtained, non-canonical bases are replaced with randomly chosen canonical bases. With a probability of 50%, the sequence is reverse complemented to ensure the model can accurately classify sequences with reverse alignments to the assembly. The errors are then introduced using BadRead [29], a long-read simulator that generates erroneous reads from perfect sequences. For BadRead, we use the ‘nanopore2020’ error model with identity parameters set to (90; 98; 5) and glitch parameters set to (10,000; 25; 25). After generating the erroneous sequence, we discard it if it is shorter than 500 bp and truncate it to 1000 bp by keeping the first 1000 bp if it exceeds this length. The resulting sequence, along with the corresponding species ID and genus ID, forms a single example used for training. Validation examples are generated in the same manner.

### 2.5 Evaluation datasets

We evaluated our model on long-read Nanopore datasets sequenced using both R9.4.1 and R10.4.1 flow cells. A list of datasets and their details is provided in Table 1.

**Table 1:** Summary of evaluation datasets used in this study.

|  | In-sample | Out-of-sample | Zymo<br>D6322 R10 | Zymo<br>D6322 R9 |
| --- | --- | --- | --- | --- |
| Num Species | 47<br>46 | 33<br>31 | 7 | 7 |
| Distribution | Even | Even | Even | Even |
| Num Reads | 141000<br>460000 | 165000<br>310000 | 140000 | 70000 |
| Min Length | 1000 | 1000 | 1000 | 1000 |
| Avg Length | 8052<br>8102 | 9811<br>9510 | 7088 | 17793 |
| Max Length | 225369 | 244499 | 66561 | 173774 |
| Flowcell | R9 | R9 | R10 | R9 |

### 2.6 Bacterial isolate datasets

The in-sample and out-of-sample datasets were introduced in [23]. These datasets consist of 81 sequenced bacterial isolates obtained from NCBI’s Sequence Read Archive (SRA), with reads sequenced using an R9.4.1 flow cell. The authors of [23] ensured that each bacterial isolate’s assembly has a matching reference genome in the MetageNN database, with an average nucleotide identity (ANI) exceeding 99%.

The 81 bacterial isolates are divided into two groups. The first group, consisting of 47 isolates, has its representative reference genomes included in the training database and is referred to as the “in-sample dataset.” The second group, containing 34 isolates, lacks representative reference genomes in the training database and is referred to as the “out-of-sample dataset.”

All but one of the species in the out-of-sample dataset have at least one closely related species from the same genus present in the training database. The single isolate without a representative from the same genus was excluded from further analysis, resulting in a total of 33 isolates in the out-of-sample dataset.

Since our model is trained on sequence chunks greater than 1000 bp, we excluded all reads from isolates with lengths shorter than 1000 bp. For read-level evaluation, we sampled 3,000 reads from each isolate in both the in-sample and out-of-sample datasets to ensure an even distribution. For community-level evaluation, we sampled 10,000 reads per isolate. However, one isolate from the in-sample dataset and two isolates from the out-of-sample dataset had fewer than 10,000 reads and were therefore discarded. As a result, the community-level evaluation included 46 isolates from the in-sample dataset and 31 isolates from the out-of-sample dataset.

### 2.7 Zymo D6322 datasets

The Zymo D6322 datasets were sequenced from the ZymoBIOMICS High Molecular Weight DNA Mock Microbial sample. Sequencing was done on GridION release 22.12.5. using SQK-NBD114.24 barcoding kit and FLO-MIN114 flowcells. Live SUP basecalling was done with Guppy 6.5.7, with default demultiplexing parameters and barcode trimming.

We subsampled 20,000 reads per species, selecting only reads with a minimum length of 1,000 bp. Since we focus on bacterial species, Saccharomyces cerevisiae reads were removed from the dataset. True labels were assigned by running minimap2 [30], retaining only high-confidence mappings with a mapping quality higher than 30 and considering only primary alignments.

The Zymo D6322 R9 dataset was previously sequenced by [31]. True labels were assigned using minimap2 in the same manner as for the R10.4.1 dataset. For this dataset, we subsampled 10,000 reads per bacterial species, applying the same 1,000 bp minimum length filter.

### 2.8 Evaluation

We evaluated our model for read-level classification and community-level abundance estimation, conducting both evaluations at the species and genus levels. For comparison with Kraken 2, we built a custom database where taxonomy information was explicitly assigned at either the species or genus level using the kraken:taxid tag [32]. This ensured a fair comparison by preventing Kraken 2 from relying on external taxonomy files. The reads that Kraken 2 failed to classify were assigned to an additional “unknown” class. Consequently, in our evaluations, these reads were always considered incorrect predictions, as this label does not exist in the ground truth.

We also compared Metaxa with MetageNN, the deep-learning tool for classifying Nanopore reads. We present MetageNN results only for genus-level classification, as the tool does not classify reads at the species level. Rafael Peres da Silva, the author of MetageNN, assisted in training and evaluating MetageNN to ensure optimal parameter selection and fair benchmarking.

### 2.9 Evaluation Metrics

For read-level classification, we computed precision, recall, F1-score, and overall accuracy. Given a class-specific confusion matrix with true positives (TP), false positives (FP), false negatives (FN), and true negatives (TN), these metrics are defined as follows:

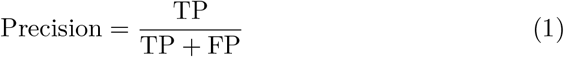

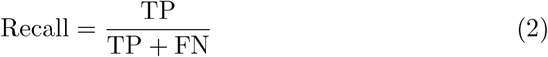

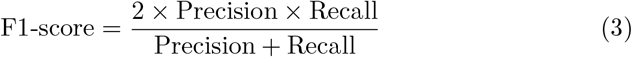

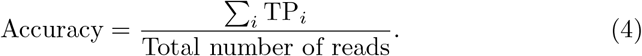

For community-level evaluation, we assessed the *L*_2_ distance between true and estimated abundances across 200 sampled datasets. If the true abundance for species *i* is *p*_*i*_, and the estimated abundance is 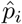, the *L*_2_ distance is defined as *L*_2_ norm between predicted abundances and true abundances:

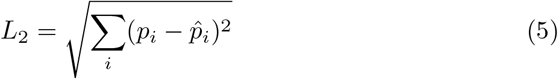

where *p*_*i*_ and 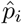 are normalized to sum to 1 across all taxa in the dataset.

Moreover, we evaluate the methods based on their precision-recall performance using average precision (AP), which summarizes the area under the precision-recall curve. The equation for average precision is given as:

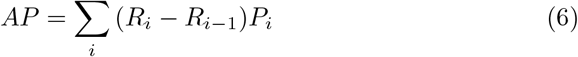

where *i* represents the abundance threshold. Species are ranked by their estimated abundances and considered present if their true abundance exceeds 10^−9^. Here, precision *P*_*i*_ refers to the proportion of predicted present species that are truly present, while recall *R*_*i*_ represents the proportion of truly present species that are correctly predicted as present.

Additionally, we evaluated classification performance at the community level based on the detection of species at abundance thresholds of 0.1%, 1%, and 10%. A species was considered detected if its estimated abundance exceeded the given threshold. We computed precision, recall, F1-score, and accuracy using the same formulas as in read-level classification, with true positives, false positives, false negatives, and true negatives defined according to species detection at each threshold.

## 3 Results

### 3.1 Read-Level Classification Performance

Table 2 shows the classification accuracy of Metaxa and Kraken 2 across different datasets at both the species and genus levels. The in-sample dataset consists of 47 species, while the out-of-sample dataset includes 33 species. The Zymo D6322 datasets, sequenced using R9 and R10 flow cells, each contain seven species. Kraken 2 outperforms Metaxa in both species- and genus-level accuracy on the in-sample dataset. However, Metaxa significantly outperforms Kraken 2 at the genus level for species that are not present in the database but have at least one closely related species from the same genus included. Compared to another deep learning-based method for genus-level classification, Metaxa significantly outperforms MetageNN on both in-sample and out-of-sample datasets. On the Zymo datasets, Kraken 2 slightly outperforms Metaxa in species-level classification, while both tools perform similarly at the genus-level. However, in genus-level classification, MetageNN lags behind both Kraken 2 and Metaxa on both Zymo datasets. Full results, including precision, recall, and F1 score for each dataset, classification level, and method, are provided in Supplementary Table 1.

**Table 2:**
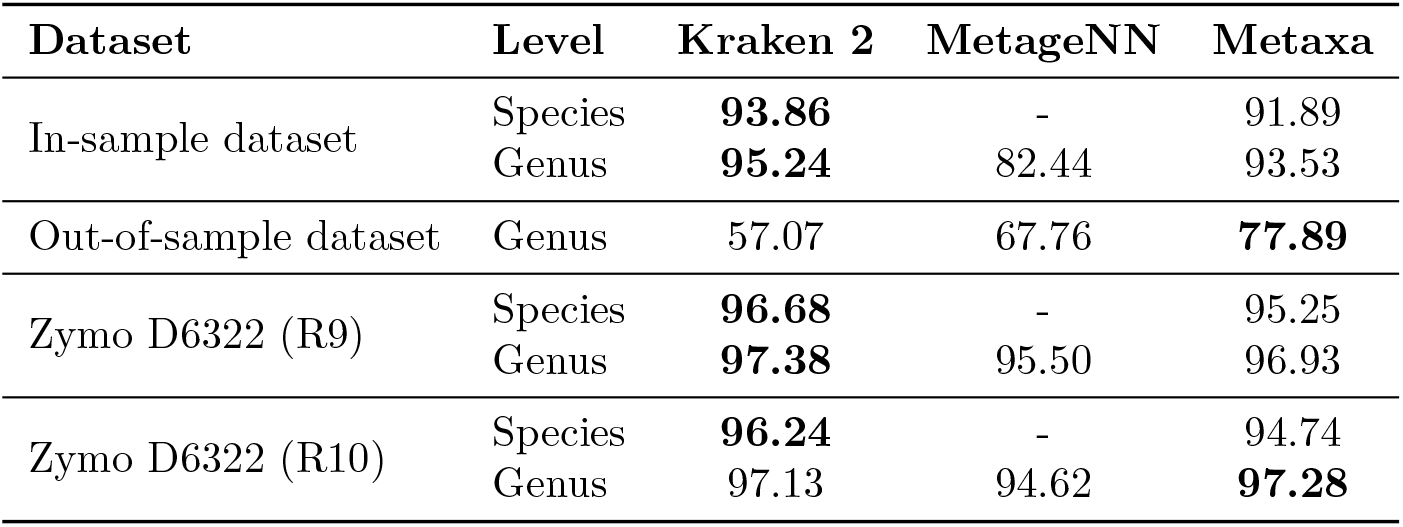
Classification accuracy (%) of the tools evaluated on different datasets at species and genus levels.

| Dataset | Level | Kraken 2 | MetageNN | Metaxa |
| --- | --- | --- | --- | --- |
| In-sample dataset | Species | **93.86** | - | 91.89 |
|  | Genus | **95.24** | 82.44 | 93.53 |
| Out-of-sample dataset | Genus | 57.07 | 67.76 | **77.89** |
| Zymo D6322 (R9) | Species | **96.68** | - | 95.25 |
|  | Genus | **97.38** | 95.50 | 96.93 |
| Zymo D6322 (R10) | Species | **96.24** | - | 94.74 |
|  | Genus | 97.13 | 94.62 | **97.28** |

Figure 2 presents the detailed read-level performance of Kraken 2 and Metaxa across different evaluation datasets and taxonomic levels. Both tools exhibit high precision, with Kraken 2 slightly outperforming Metaxa across all datasets and taxonomic levels. Additionally, both tools achieve high recall, except for the out-of-sample dataset. For this dataset, Metaxa attains significantly higher recall than Kraken 2 and, consequently, a higher F1 score. Furthermore, both methods share most outliers, suggesting that lower performance for certain species or genera is more likely due to external factors, such as dataset quality or reference assembly, rather than biases inherent to either method. Detailed precision, recall, and F1 scores for each species and genus, across datasets and methods are provided in Supplementary Tables 2–5.

**Figure 2:**
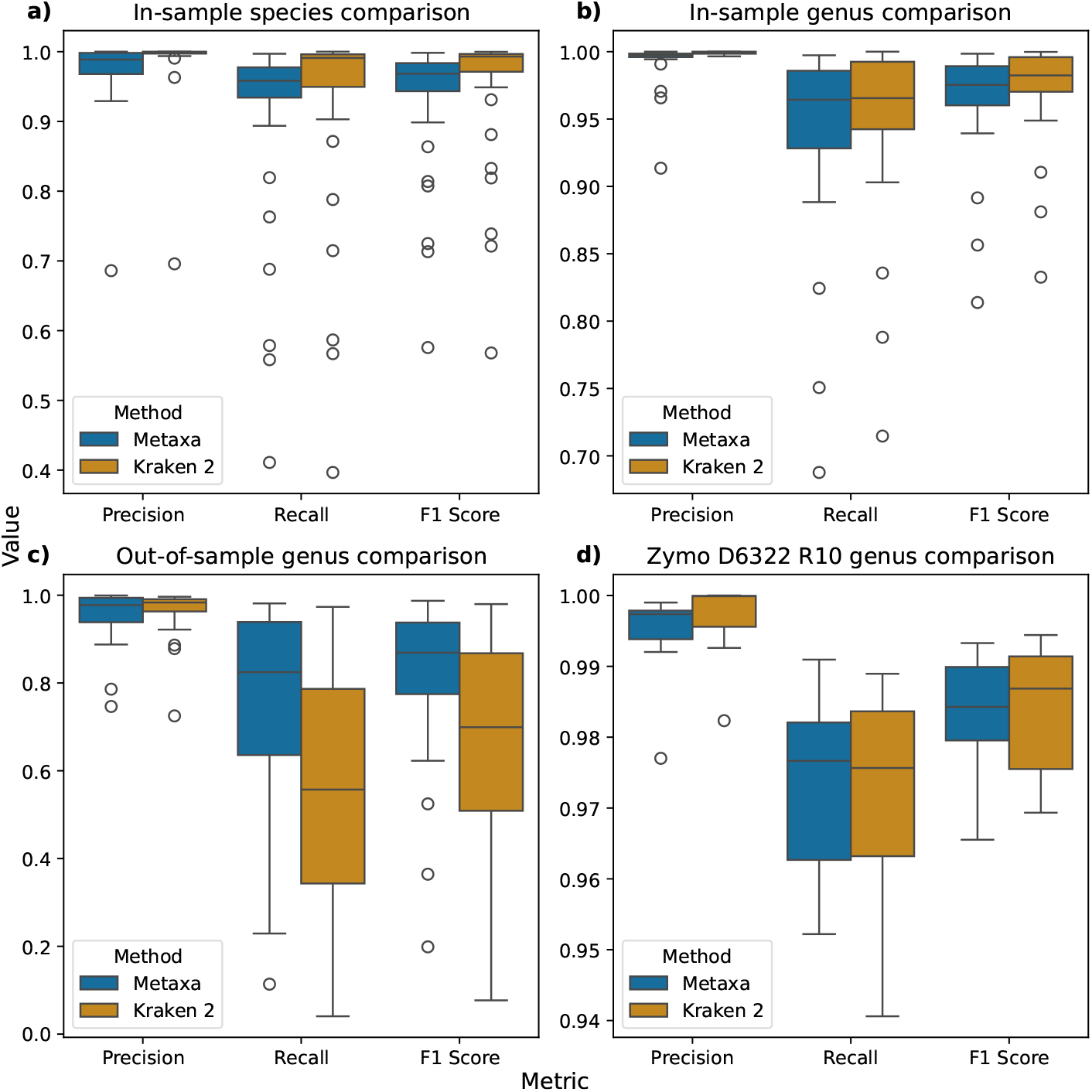
Read-level classification performance of Metaxa and Kraken 2 across different datasets and taxonomic levels, presented as boxplots for precision, recall, and F1 score. Subfigure (a) shows species-level classification performance on the in-sample dataset, while subfigure (b) presents genus-level classification performance on the same dataset. Subfigure (c) illustrates genus-level classification performance on the out-of-sample dataset, and subfigure (d) depicts genus-level classification performance on the Zymo D6322 dataset. In each box plot, the median is represented by the central line, whiskers extend to 1.5 times the interquartile range (IQR), and points outside this range are considered outliers.

### 3.2 Community-Level Evaluation

We evaluated community-level classification performance using mock microbial communities. Each mock community was generated by randomly selecting 10 species (or 10 genera) from the dataset and subsampling the reads according to a log-normal distribution with parameters *µ* = 3 and *σ* = 1.5. This process was repeated for *N* = 200 iterations to assess the variability in classification accuracy.

For each iteration, we calculated the *L*_2_ distance between the true and estimated community composition. Additionally, classification accuracy, precision, recall, and F1 score were calculated at detection thresholds of 0.1%, 1%, and 10%.

Figure 3 compares the community-level performance of Kraken 2 and Metaxa. Both methods exhibit similar *L*_2_ distance (Figure 3a) and average precision between predicted and true abundances (Figure 3b). The most noticeable difference is in the F1 score for low-abundance species (threshold 0.001; see Figure 3c), where Kraken 2 outperforms Metaxa. However, at higher abundance thresholds, their scores converge. Another key difference is Kraken 2’s failure to detect high-abundance taxa (threshold 0.1) at the genus level for out-of-sample data, whereas Metaxa’s score at this threshold exceeds its performance at lower thresholds, consistent with its species-level classification performance on in-sample data Figure 3d). The complete results for each experiment on both the in-sample and out-of-sample datasets are provided in Supplementary Tables 6–9.

**Figure 3:**
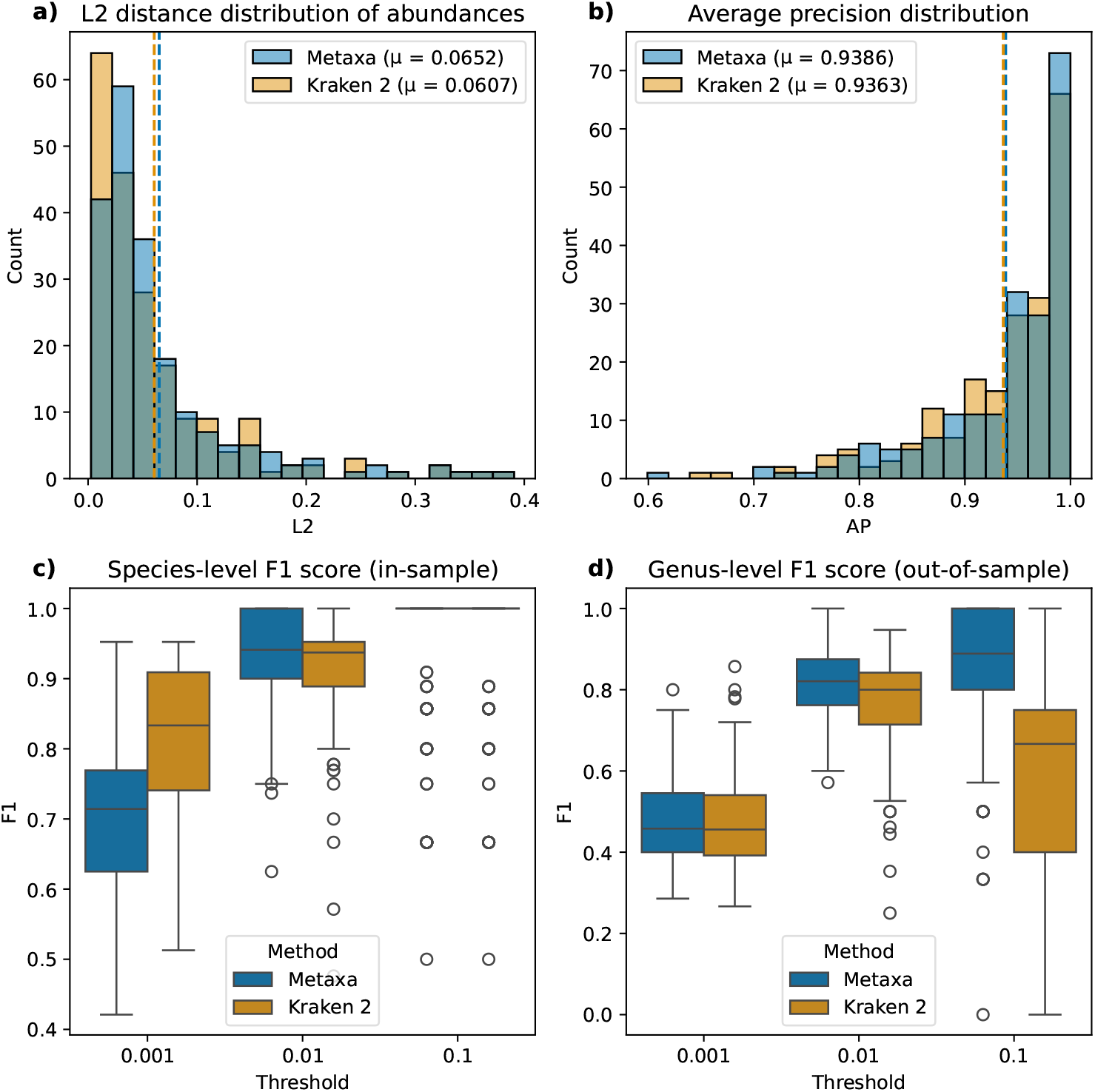
Community-level classification performance of Metaxa and Kraken 2. Subfigure (a) shows the distribution of L2 distances between predicted and true relative abundances. Subfigure (b) presents the distribution of average precision (AP) values for both methods. Subfigure (c) illustrates F1 score-based classification performance at different abundance thresholds for species-level classification in the in-sample dataset, while subfigure (d) depicts F1 score-based classification performance at different abundance thresholds for genus-level classification in the out-of-sample dataset. For box plots, the central line represents the median, whiskers extend to 1.5 times the interquartile range (IQR), and points beyond this range are considered outliers.

### 3.3 Effects of length and quality on accuracy

Lastly, we evaluated the classification performance of Metaxa with respect to read quality and read length. To do this, we generated a total of 1 Gbp of simulated reads using BadRead from the training database, with length parameters set to 2000, 1000 and read identity parameters set to 95, 100, 5. After generating the reads, we removed those shorter than 1 kbp as well as reads classified as chimeric, junk, or random. Figure 4 illustrates the effects of read length and read quality on classification accuracy at both species and genus levels. As expected, the classification accuracy improves with increasing read identity and, in most cases, with increasing read length. This correlation arises because a higher read identity corresponds to fewer sequencing errors, resulting in sequences that are more similar to those in the training database. Additionally, since classification is performed on fixed-length chunks, longer reads produce more chunks, leading to a greater number of predictions available for aggregation. Species- and genus-level classification accuracies for each read identity and read length bin are provided in Supplementary Tables 10–11.

**Figure 4:**
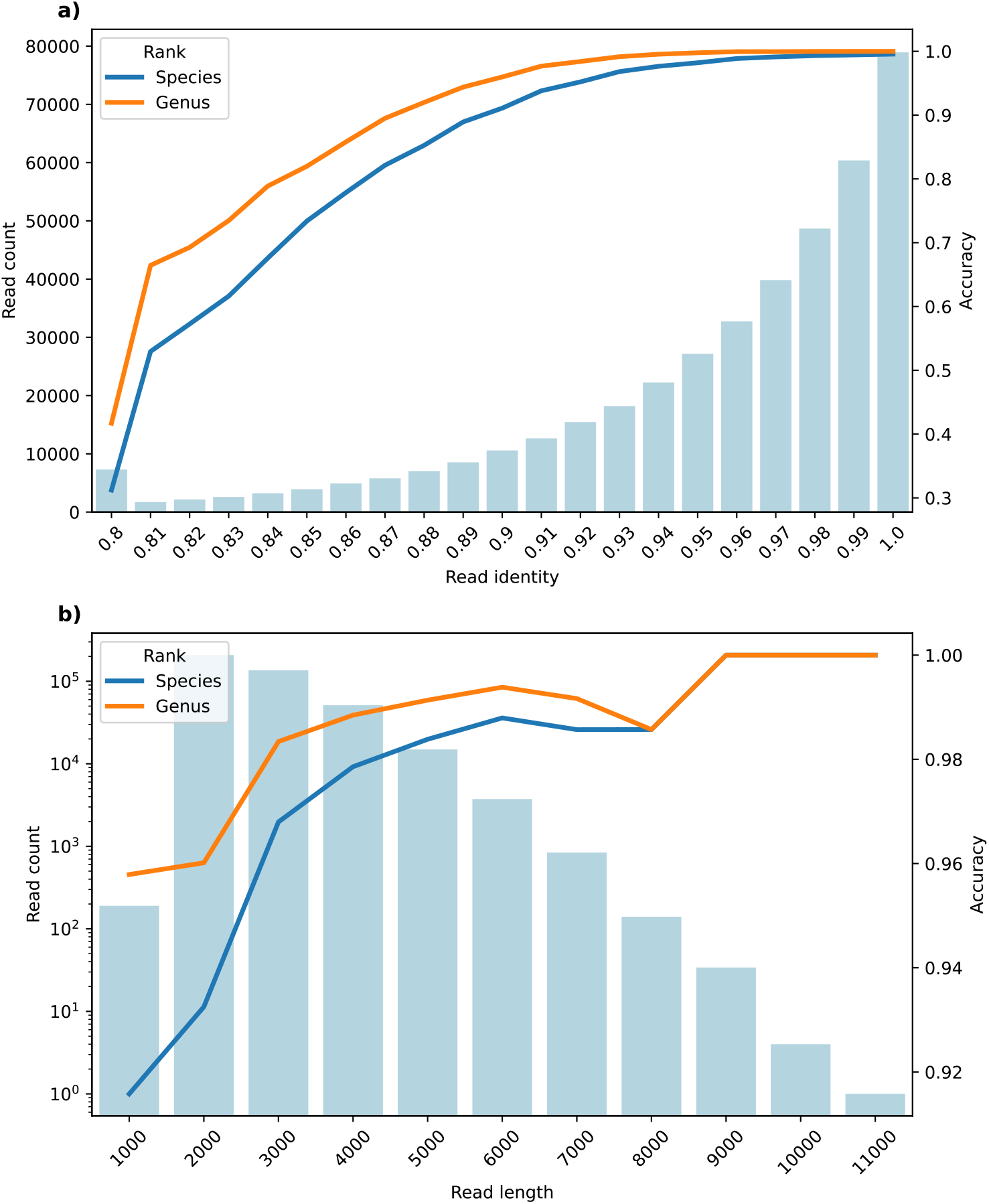
Classification accuracy of Metaxa at the species and genus levels as a function of read identity (a) and read length (b). Read identity is binned from 0.8 to 1.0 in increments of 0.01, with all values below 0.8 grouped into the 0.8 bin. Read length is binned from 1000 bp to 11,000 bp in 1000 bp increments. Each bin includes values up to and including the right boundary (e.g., the 2000 bp bin includes reads up to 2000 bp). The histograms represent the number of reads in each respective bin. The figure shows that classification accuracy improves with increasing read identity and, in most cases, with longer read length.

## 4 Discussion

In this paper, we introduce Metaxa, a transformer-based tool for taxonomic classification in metagenomics. By leveraging a deep-learning architecture, Metaxa effectively captures the rich information present in DNA sequences beyond fixed k-mer representations, enabling accurate classification. Our validation experiments show that Metaxa outperforms MetageNN and performs comparably to Kraken 2 when classifying genomes included in the training database, both at the species and genus levels. Notably, Metaxa significantly outperforms Kraken 2 and MetageNN at the genus level for species absent from the training data, highlighting its ability to generalize by capturing sequence context over longer regions rather than relying solely on individual k-mers or k-mer profiles. This capability is particularly valuable for analyzing novel species in microbiome studies, pathogen detection, and environmental metagenomics, where complete reference genomes are often unavailable or incomplete.

In our comparative analysis, we selected Kraken 2 as the baseline for evaluating Metaxa. Kraken 2 is a widely adopted k-mer-based taxonomic classification tool known for its high accuracy and efficiency in processing metagenomic data. While there are other k-mer-based approaches, other methodologies such as read mapping approaches like minimap2, and protein sequence matching tools such as Kaiju [33], Kraken 2’s prominence and performance make it a suitable representative of classical algorithm-based tools.

Our objective was to demonstrate the efficacy of deep learning models in taxonomic classification tasks. Notably, our findings indicate that deep learning tools can achieve performance on par with state-of-the-art tools like Kraken 2 when all reference sequences are present in the database. Moreover, Metaxa performs better in scenarios where certain references are absent, highlighting its robustness and adaptability in handling incomplete databases or previously unknown microbial taxa.

The potential of deep learning in metagenomic classification is evident, as shown by Metaxa’s species-level performance, which matches Kraken 2. Unlike traditional heuristic-based methods, deep-learning models can more effectively capture complex sequence patterns and relationships across long genomic regions. As these architectures continue to evolve in bioinformatics, advancements in accuracy, interpretability, and computational efficiency could establish them as the preferred approach for taxonomic classification.

Despite these promising results, Metaxa has several limitations. Its training dataset is limited in size, as we used only a small subset of reference genomes from the RefSeq database, which may impact its taxonomic coverage. Expanding the training dataset to include a broader and more diverse set of reference genomes could further improve Metaxa’s ability to classify novel species accurately. Additionally, training is computationally expensive compared to constructing a Kraken 2 database. Future work should focus on improving training efficiency while maintaining classification accuracy, potentially by optimizing the training pipeline or exploring more efficient model architectures. Techniques such as model pruning, distillation, or dataset compression could be explored to reduce inference time and make Metaxa more scalable for large metagenomic datasets.

Another limitation arises from our training approach, which utilizes only a single representative genome per species. A more comprehensive dataset, including multiple strains and real-world metagenomic samples, could improve performance and robustness. Similarly, while we use BadRead for error simulation, its synthetic profiles may not fully capture the complexity of real Nanopore sequencing errors. Future efforts could explore refined error models or train directly on real Nanopore reads.

Metaxa demonstrates the power of deep learning in metagenomic classification, particularly for genus-level assignments of species without a reference genome in the database. By leveraging a transformer-based architecture, Metaxa significantly outperforms both MetageNN and Kraken 2 in these challenging cases, where only a related species within the same genus is available. This highlights its ability to generalize beyond exact reference matches, making it a valuable tool for analyzing novel or underrepresented taxa. Additionally, a key strength of Metaxa is its use of BadRead to simulate Nanopore sequencing errors during training. Since it is not feasible to obtain real Nanopore reads for every reference genome in the database, and training directly on high-quality assemblies is suboptimal due to their disparity from Nanopore reads, this approach ensures a more realistic training process. Furthermore, Metaxa’s performance, which is on par with Kraken 2, highlights the potential of deep-learning approaches for species-level classification with ONT data. As long-read sequencing technologies continue to evolve, deep-learning models like Metaxa are poised to become increasingly effective, bridging the gap between raw sequencing data and high-accuracy taxonomic classification.

## 5 Data availability

The source code for Metaxa, along with metadata describing the training database, is available at https://github.com/lbcb-sci/metaxa or at Zenodo https://zenodo.org/records/15309015. The trained model can be accessed via Zenodo at https://zenodo.org/records/15062544.

The SRA accession lists used to construct the in-sample and out-of-sample datasets were originally compiled by the MetageNN authors [23] and are available on their GitHub repository at https://github.com/CSB5/MetageNN/tree/d7c858cee1b88bf78fb1ea7d120b3be180a52b69

Public Zymo D6322 R9 Nanopore sequencing data were obtained from the UNCALLED study [31], available under NCBI BioProject PRJNA604456. Newly sequenced Zymo D6322 R10 data are available under BioProject PRJNA1240873.

## 6 Acknowledgements

We thank Rafael Peres da Silva for his assistance in training and evaluating MetageNN on the datasets described in this study. We also thank Martin Frederik Schmitz, who helped us draw the graphic abstract.

## Author contributions

M.S. conceived the study; K.F., D.S. and M.S. designed the research; D.S. and K.F. performed the research; D.S. designed and implemented the model; B.C. sequenced the dataset; D.S. and K.F. drafted the manuscript. All authors read and approved the final version of the manuscript.

